# Validation of an oral self-administration model of xylazine use in mice

**DOI:** 10.64898/2026.06.02.729589

**Authors:** Reece C. Budinich, Lars H. Nelson, Max E. Joffe

**Affiliations:** Department of Psychiatry, University of Pittsburgh, Pittsburgh PA; Translational Neuroscience Program, University of Pittsburgh; Center for Neuroscience at the University of Pittsburgh

**Author notes:** Max Joffe, 219 BDG2, 450 Technology Drive, Pittsburgh, PA 15219.

## Abstract

Xylazine is a pervasive adulterant in clandestine opioid supplies. This trend is troubling, as xylazine carries its own acute side effects and its long-term cognitive and motivational effects are not known. Thus, we developed and validated a model of oral xylazine self-administration that is conducive to polysubstance studies and can easily be implemented in animal research labs. Mice underwent 4-hour drinking sessions where their only source of drinking water was adulterated with xylazine (10-1000µg/mL). Oral bioavailability and brain penetrance were validated using mass spectrometry on brain and plasma samples collected after a drinking session. Male and female mice decreased fluid intake at high concentrations of xylazine, with male mice consuming less than female mice at intermediate concentrations (100 and 300µg/mL). Female mice had decreased locomotion following drinking sessions at 100, 300, and 1000µg/mL. After a 4-hour drinking session, all mice received brain xylazine concentrations that are above the Ki (affinity) and EC50 (potency) of several known binding targets of xylazine. We then performed a three-bottle choice experiment where mice had the option of drinking from water with xylazine, water with fentanyl, or plain water. In three-bottle choice, mice consumed less xylazine-containing water than fentanyl-containing or plain water. Our results indicate that xylazine is orally bioavailable in mice, that mice will readily drink xylazine to pharmacologically and behaviorally relevant doses, and that mice prefer consuming fentanyl and water over xylazine.

## Main

The veterinary sedative xylazine has become a widespread adulterant in clandestine opioids. This adulteration is commonly attributed to xylazine’s ability to cause similar sedation to opioids and/or prolong fentanyl’s short duration of action. People who use illicit opioids generally dislike xylazine; consistent with this, preclinical studies have found that xylazine decreases opioid-seeking and reward-associated behaviors (1). Nonetheless, the preclinical literature is scarce and sometimes conflicting, and the long-term cognitive and motivational effects of xylazine remain unclear.

Polysubstance use, like xylazine-opioid combinations, is prevalent, but relevant preclinical research is technically challenging (2). While intravenous self-administration of drugs is considered the gold-standard preclinical model to assess reinforcing properties of drugs (3), several challenges limit its widespread use in rodents. First, the procedures are technically demanding and time-intensive, and challenges become magnified when additional surgeries (e.g., intracranial cannulation) are required. Second, the small vasculature of mice generally limits the length of experiments to 2- weeks or less, hindering studies on long-term effects of drug exposure. Finally, dual jugular cannulation is needed to assess simultaneous reinforcing properties of two drugs, but this laborious technique is rare in rats and non-existent in mice in the current literature. To meet these challenges, we set out to develop and validate a means of xylazine self-administration that is widely accessible and conducive to assessing complexities inherent in polysubstance research.

Oral self-administration is used in rodent models to study voluntary intake of alcohol, nicotine, opioids, and other drugs. We therefore aimed to validate xylazine’s oral bioavailability in mice and determine relevant drinking concentrations. We first performed a concentration-response experiment where mice only had access to water adulterated with xylazine (single-bottle-choice) (Figure 1A-1G). For 4 hours per day, mice were placed into cages where all water was provided by a single sipper bottle (Figure 1A). The concentration of xylazine in water increased from 0 – 1000 µg/mL every 2 days. On every second day, mice underwent an open field locomotor activity assessment immediately following the drinking session. As the xylazine concentration increased, mice decreased fluid consumption (Figure 1B). Female mice drank more fluid (Figure 1B) and consumed more xylazine (Figure 1C) than males at 100-300 µg/mL. Additionally, females displayed reduced locomotion following access to 100- 1000 µg/mL xylazine (Figure 1D), consistent with sedation. By contrast, male locomotion was unaffected at these doses. There was a strong negative correlation between xylazine consumption and locomotion at the 1000 µg/mL dose (R^2^=0.6961; p=0.0014) (Figure 1E), suggesting the sex difference in sedation was driven by dose.

**Figure 1.**
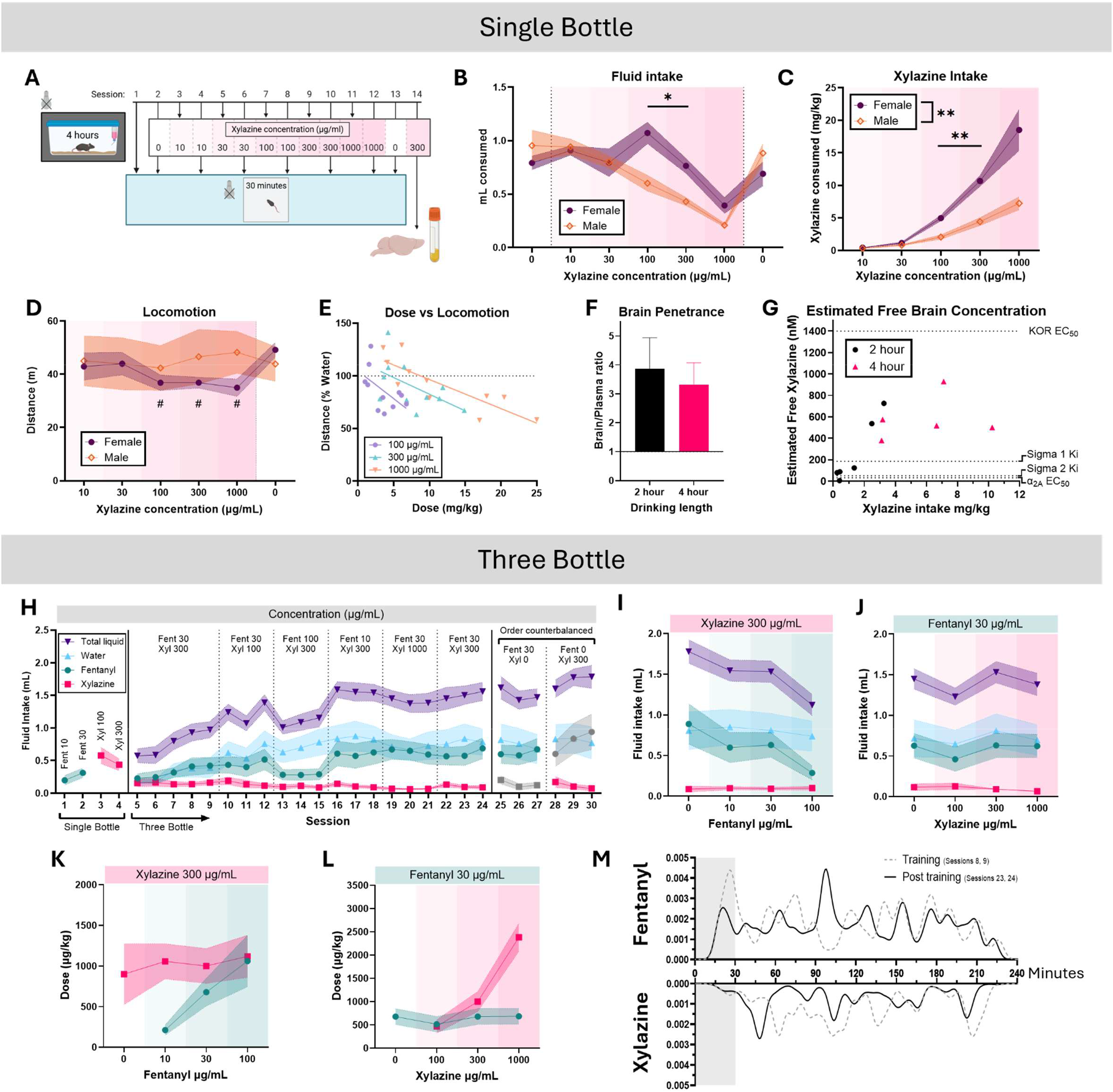
Xylazine oral administration in mice. (A) Timeline of single-bottle-choice concentration-response. **(B)** Volume consumed (concentration-sex interaction F_(3.229, 28.52)_=8.788, ***p =0.0002; Šídák’s multiple-comparisons test Male vs Female 100µg/mL: *p=0.0429; 300µg/mL: *p=0.0379), **(C)** dose consumed (concentration-sex interaction F_(1.088, 9.795)_=12.57, **p=0.0049; multiple-comparisons test Male vs Female 100µg/mL:**p = 0.0040; 300µg/mL:**p=0.0019), **(D)** and distance traveled (concentration-sex interaction F_(5, 45)_=3.313, *p=0.0124; multiple-comparisons test vs Female 0µg/mL 100µg/mL:#p=0.0378; 300µg/mL:#p=0.0395; 1000µg/mL:#p=0.0095) after drinking in single-bottle experiment. **(E)** Correlations between xylazine dose consumed and reduced locomotion (relative to plain water). **(F)** Brain penetrance of orally-consumed xylazine. **(G)** Estimated xylazine free brain concentration vs dose intake after oral administration. Horizontal lines represent xylazine’s Ki at sigma receptors and EC_50_ at α2A receptors and kappa opioid receptors (affinity/potency values from (4)). **(H)** Three-bottle-choice timeline and fluid consumption per bottle per session. **(I)** Fentanyl and **(J)** xylazine concentration-fluid intake response curve. **(K)** Fentanyl and **(L)** xylazine concentration-dose response curves. **(M)** Sipper interaction time-course density analysis for sessions 8/9 (training) and 23/24 (post training). Shaded region highlights the first 30 minutes. Made with biorender.com (https://BioRender.com/6wxm4ce).

We next performed pharmacokinetics studies to confirm oral bioavailability of xylazine. Equilibrium dialysis revealed that ∼36% of total xylazine incubated in mouse brain tissue is free/unbound (Figure S1A). Next, we collected plasma and brain tissue from mice after oral administration of 300 µg/mL and used mass spectrometry to determine levels of xylazine and common metabolites (Figure S1B). Mice were sacrificed after 2- or 4-hours access. The total brain/plasma ratio was 3.56, confirming xylazine is centrally penetrant and accumulates in brain after oral consumption (Figure 1F). Furthermore, estimated free brain xylazine concentrations reached 379-928 nM after 4 hours of xylazine drinking (Figure 1G). All animals that consumed more than 3 mg/kg xylazine achieved estimated free brain concentrations sufficient to engage sigma receptors and α2 adrenergic receptors (4). Collectively, these data indicate that mice can consume behaviorally and pharmacologically relevant doses of xylazine from oral administration.

To explore whether mice will voluntarily drink xylazine and how this choice may affect fentanyl intake, we instituted a 3-bottle choice design (Figure 1H-M). To acclimate mice to xylazine and fentanyl, single-bottle access was provided to either substance for the first 4 sessions. Next, for 4 hours per day, mice received access to three sipper bottles, containing (1) water with xylazine, (2) water with fentanyl, and (3) plain water. Early in training, mice quickly started to avoid drinking xylazine and increased drinking fentanyl and water (Figure 1H). We next varied the concentration of either substance for 3 consecutive days (xylazine=0-1000µg/mL or fentanyl=0-100µg/mL). Mice continued to avoid xylazine and titrated the amount of fluid consumed across various concentrations of xylazine or fentanyl (Figure 1I-J). Varying the fentanyl concentration affected the fentanyl but not xylazine dose, while changing the xylazine concentration affected xylazine intake but not fentanyl (Figure 1K-L). However, intake of xylazine was at such low levels (max ∼ 2.5 mg/kg/session) that mice may not have consumed sufficient xylazine to alter fentanyl directed behavior. We also monitored visits to each sipper over several sessions and found that mice were more likely to interact with the fentanyl sipper earlier in the session than the xylazine sipper (Figure 1M).

In summary, we found that xylazine is orally bioavailable in mice, and that mice can be made to orally self-administer xylazine to pharmacologically and behaviorally relevant brain concentrations. These findings support the validity of oral administration as a model of xylazine use and inclusion of xylazine in models of fentanyl oral self-administration. Furthermore, we found that mice prefer consuming fentanyl or plain water over xylazine, consistent with reports of undesirability in humans. Future studies may leverage the pharmacokinetics and dose-response relationships established here to design translational experiments assessing xylazine’s reinforcing properties and long-term effects in mice.

## Funding Support

This work was supported by. This work is the result of NIH funding (PI: Joffe DP1DA060482, R01DA058704, R21DA062048; PI: Gelhaus NIHS10OD032141), in whole or in part, and is subject to the NIH Public Access Policy. Through acceptance of this federal funding, the NIH has been given a right to make the work publicly available in PubMed Central.

## Methods

### Animals

All animals were maintained in reverse light cycle (7am lights off, 7pm lights on) with ad libitum access to food and water. All experimentation occurred > 2.5 hours into the dark phase.

### Drugs

Xylazine hydrochloride (PHR3264) and fentanyl citrate (PHR8977) were purchased from Sigma-Aldrich and dissolved in tap water.

### Single-bottle xylazine administration

#### Habituation

Mice (C57BL/6J; n = 6 male, 5 female; 11-14 weeks old at start of experiment) were handled for >1 minute on day one. On day two, mice were placed in a 30 × 30 cm plexiglass open field chambers for 30 minutes. On day three, mice were removed from their home cages and individually placed into new cages where their water was provided by glass sipper bottles. After 4 hours, the sipper bottles were removed and mice were transferred to an experimental room outside the housing facility and placed in open field chambers for a 30-minute locomotion session (transit from bottles removed to starting locomotion recording = ∼6 minutes) where distance traveled was measured using ANY-maze software. Sipper bottles were weighed before and after this 4-hour session to measure liquid consumption. After the 30-minute locomotion session, mice were returned to their home cages in the housing room.

#### Concentration response

Xylazine was added to the sipper bottle in an ascending concentration order (10, 30, 100, 300, 1000 µg/mL) with 2 days for each concentration. For the first day of each concentration, the mice had a 4-hour drinking session without locomotion testing. On the second day of each concentration, mice had both the 4-hour drinking session and 30-minute locomotion testing (as described above). Following the 1000 µg/mL concentration, a 0 µg/mL (i.e., only tap water) drinking session with locomotion testing was performed.

### Xylazine brain and plasma concentrations following drinking

Mice from the single-bottle concentration-response experiment were used to determine brain and plasma concentrations after xylazine drinking. Mice were randomly assigned to either a 2-hour (n = 3 male, 3 female) or 4-hour (n = 3 male, 2 female) drinking session at 300 µg/mL. Bottles for the 4-hour group were measured both at 2 hours and 4 hours. Immediately after the session ended, mice were briefly anaesthetized with isoflurane and decapitated. Trunk blood was collected in heparinized tubes (Microvette CB300) and was centrifuged at 2000 RPM for 15 minutes at 4°C. Plasma was then pipetted into a new tube. The brain was extracted and frozen on dry ice. Plasma and brains were stored at -80°C until mass spectrometry.

### Xylazine-brain binding – equilibrium dialysis

#### Brain homogenate preparation

Brains from 4 experimentally naïve mice (n = 2 male, 2 female; 22 weeks old) were extracted on ice and weighed. Brains were combined with 1X PBS (1:3 w/v brain:PBS) in a 15mL tube and homogenized by ultrasonic probe [on ice]. Two aliquots of brain homogenate were made, one of which was spiked with xylazine to a final concentration of 1.5 µM (the average total concentration of the 4-hour drinking group from the oral bioavailability experiment).

#### Rapid equilibrium dialysis

Equilibrium dialysis was performed with six technical replicates. In each replicate, 300µL of spiked brain homogenate was added to the sample/donor chamber of a rapid equilibrium dialysis (RED) device (Thermo Scientific; cat. # 90006). Each corresponding buffer/receiver chamber received 500µL of 1X PBS. The RED device was covered with microplate sealing tape and placed in an incubator/shaker set to 37°C and 250 RPM for 16 hours. In duplicate, 100µL was taken from the red chamber (spiked tissue homogenate) and 100µL was taken from the white chamber (PBS buffer) and added to separate microcentrifuge tubes. To match the total biological content of each sample, 100µL 1X PBS was added to the tubes containing dialyzed brain homogenate, and 100µL blank brain homogenate to the tubes containing dialyzed PBS. Samples were then stored at -80°C until mass spectrometry.

### Untargeted high-resolution LC-HRMS

#### Sample preparation

Metabolic quenching and polar metabolite pool extraction was performed by adding ice cold PBS (at a ratio of 1:15 wt input tissue:vol. Deuterated (D3)-creatinine and (D3)- alanine, (D4)-taurine and (D3)-lactate (Sigma-Aldrich) was added to the sample lysates as an internal standard for a final concentration of 10µM. Samples are homogenized using a MP Bio FastPrep system using Matrix D (ceramic sphere) for 60 seconds at 60hz. 50µL lysate was transferred to a clean Eppendorf containing 200µL methanol. After vortexing, the supernatant was then cleared of protein by centrifugation at 16,000xg. 200µL cleared supernatant was dried to completion under N2 gas and resuspended in 40µL water/0.1% formic acid. 2µL of cleared supernatant was subjected to online LC-MS analysis.

#### LC-HRMS Method

Analyses were performed by untargeted LC-HRMS. Briefly, Samples were injected via a Thermo Vanquish UHPLC and separated over a reversed phase Phenomenex Kinetex C18 column (2.1×150mm, 1.7μm particle size) maintained at 55°C. For the 15 minute LC gradient, the mobile phase consisted of the following: solvent A (water / 0.1% FA) and solvent B (ACN / 0.1% FA). Flow rate is 300µL/min. The gradient was the following: 0-1min 5% B, increase to 15%B over 4 minutes, continue increasing to 98%B over 5 minutes, hold at 98%B for five minutes, reequillibrate at 1%B for five minutes. The Thermo Exploris 240 mass spectrometer was operated in positive ionization mode, scanning in ddMS2 mode (2 μscans) from 100 to 800 m/z at 120,000 resolution with an AGC target of 2e5 for full scan, 2.5e4 for ms2 scans using HCD fragmentation at stepped 20,35,50 collision energies. Source ionization setting was +3.5kV spray voltage, source gas parameters were 45 sheath gas, 12 auxiliary gas at 320°C, and 3 sweep gas. Vaorizer temperature was 350°C. Calibration was performed prior to analysis using the PierceTM FlexMix Ion Calibration Solutions (Thermo Fisher Scientific). Integrated peak areas were then extracted manually using Quan Browser (Thermo Fisher Xcalibur ver. 2.7) before normalization to internal standards for relative quantification. Purified standards were then purchased and compared in retention time, m/z, along with ms2 fragmentation patterns to validate the identity of compounds.

### Xylazine-Fentanyl-Water Three Bottle Choice

#### Habituation

Mice (C57BL/6J; n = 6 male, 6 female; 18-21 weeks old at start of experiment) were habituated to experimenter handling for >1 minute on the day prior to the experiment beginning. Mice were removed from their home cages and placed single-housed into new cages containing a 3-bottle sipper holder with capabilities of measuring sipper interactions with 2 of the bottles using infrared beam breaks (Figure S1C), modified from (5). Experimental cages for 3-bottle choice had their bedding removed (but cotton enrichment square present) to enable infrared beam sipper tracking, as bedding would frequently break the beam. For the first session, only a single bottle was present, which contained 10 ug/mL fentanyl. After 4 hours, the mice were removed from the experimental cage and returned to their home cages. Sipper bottles were weighed before and after this 4-hour session to measure liquid consumption. The following session had this bottle containing 30 µg/mL fentanyl. For the next two sessions, the fentanyl bottle was not present, and a bottle containing xylazine (100 µg/mL and 300 µg/mL for sessions 3 and 4 respectively) was in a different position. The left and right assignment of fentanyl/xylazine was counterbalanced between mice, but position within mice remained consistent throughout the experiment to support discrimination.

#### Three-bottle choice

For sessions 5-9 the sipper holders contained 3 bottles – one with xylazine (300 µg/mL), one with fentanyl (30 µg/mL), and one with water (tap).

#### Sipper tracking

To clean the sipper tracker data, we removed bouts longer than 30 seconds. These bouts comprised 1.05% of all bouts for the entire dataset including data that was not included in the publication. The removed bouts had a median length of 106 seconds, a mean of 206 seconds, and a maximum of 2206 seconds. We attributed these long bouts to something besides a mouse activating the sipper (e.g., bedding blocking the infrared detector). Additionally, we noticed periods where both sippers were activated. These events were not screened out. To create the sipper density plots, sipper activation periods were binarized then convolved with a gaussian kernel with a sigma of 24,000 (240 seconds) to smooth the data. The last two days of training (sessions 8 and 9) were averaged and reported as “training,” and the last two days of the concentration response where fentanyl and xylazine were returned to training concentrations (sessions 23 and 24) where averaged and reported as “post training.”

### Statistics

Statistics were performed using GraphPad Prism 10. Mixed effects analysis (Figure 1B) or repeated measures two-way ANOVA (Figures 1C-D) were used followed by Šídák’s multiple comparisons test. A simple linear regression was used to assess correlation for data in Figure 1E. Results were considered significant when p<0.05.

## Notes

### Competing Interest Statement

The authors have declared no competing interest.

