## Supplemental Data for "Validation of an oral self-administration model of xylazine use in mice"

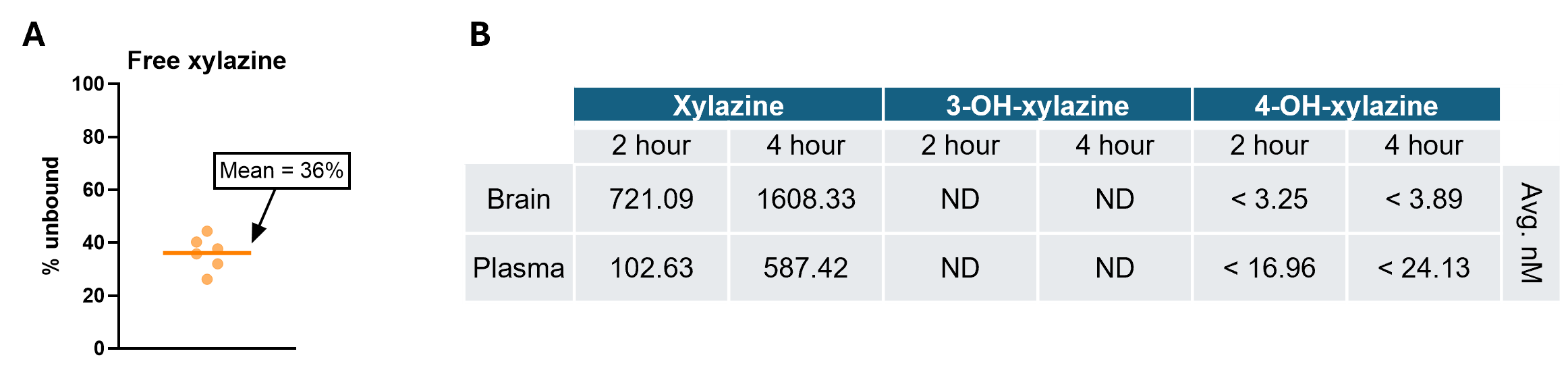

**Figure S1. (A)** 36% of total brain concentration of xylazine is unbound from tissue. **(B)** Average brain and plasma concentration of xylazine, 3-OH-xylazine, and 4-OH-xylazine after 2- or 4-hour 300µg/mL xylazine drinking from samples where detected. Xylazine was detected in all samples, 3-OH-xylazine was not detected in any samples, 4-OH-xylazine was detected in 1/6 2-hour brain samples, 2/5 4-hour brain samples, 1/5 2-hour plasma samples, and 4/5 4-hour plasma samples.

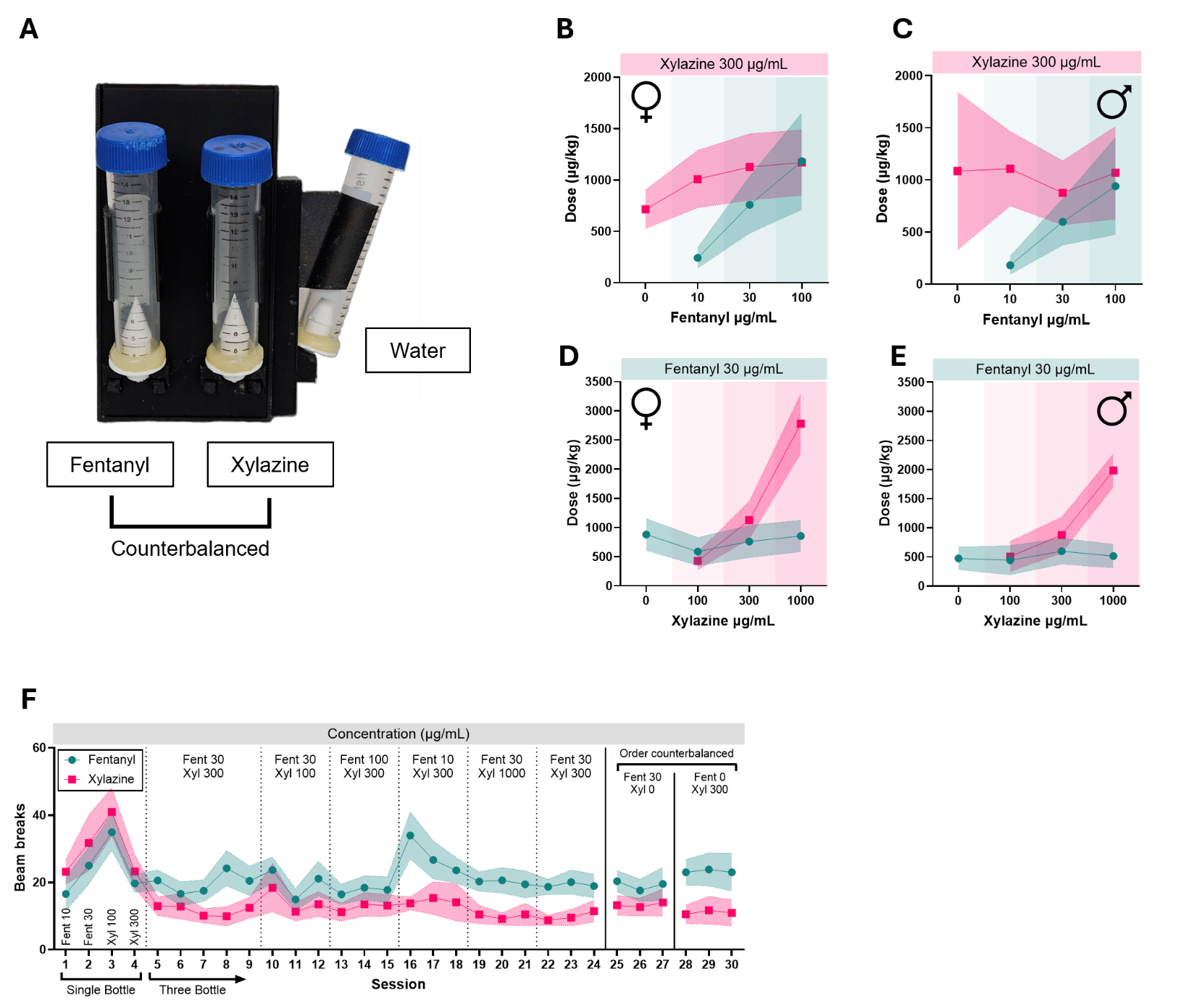

**Figure S2.** (A) Three-bottle choice sipper holder modified from Godynyuk et al. (2019) to hold a third bottle (water; right side). (B) Female and (C) male fentanyl concentration-dose consumed response curve. (D) Female and (E) male xylazine concentration-dose consumed response curve. (F) Fentanyl and xylazine sipper beam break count for three-bottle choice experiment.
